# Generating the sigmoid form in neuroscience: A threshold-crossing model

**DOI:** 10.1101/2023.11.27.568902

**Authors:** Andrew R Auty

## Abstract

The conventional interpretation^1^ of a logistic curve in biology is that it indicates the presence of an infection-like mechanism bounded by a saturation limit e.g. saturation occurs when the whole population has been infected. However, a logistic form may be generated by the fluctuating approach of biological systems to, and then exceeding, an effect threshold.

The commonly observed trends in frequency and degree of cognitive loss in neurodegenerative disease (NDD) are consistent with a threshold-crossing model. Whereas direct evidence of real life NDD being driven by an infection-like mechanism remains elusive^2,3^.

Variation in susceptibility to NDD may be explained by variation in the rate of biological ageing (e.g. neurodegeneration rate, or, metabolic interference), height of threshold and initial reserve. Each is potentially modifiable and clinically informative.

## 1. Introduction

It is believed that neurodegeneration spreads, infection-like, from brain part to brain part^4^, the infection-like agent being some kind of self-replicating protein structure, operating in the style of prion disease. Such protein structures can be observed in people who later develop symptoms of NDD, and in people who don’t. Risk of developing NDD symptoms is higher in people with these protein structures.

Clinical progress of many kinds of neurodegenerative disease (NDD) is characterised by a gradually widening scope of damaged brain parts, by the increased frequency of symptom events and by accelerated cognitive decline as the patient advances towards diagnosis^5,6^. Given that the cause of loss of function is the loss of neurones it is worth considering the possibility that the underlying neurone loss both spreads and accelerates, i.e., possess classic features of an infection-like mechanism.

Theoretical neuroscience often turns to the mathematics of infection to describe these clinical phenomena. In particular the mathematical form known as the logistic^2^ (Fig.1). This is a useful description of infection spread when the total number of initially uninfected targets is fixed. If correct, the model should have predictive power and so enable early detection of the benefits of medical interventions.

**Fig. 1.**
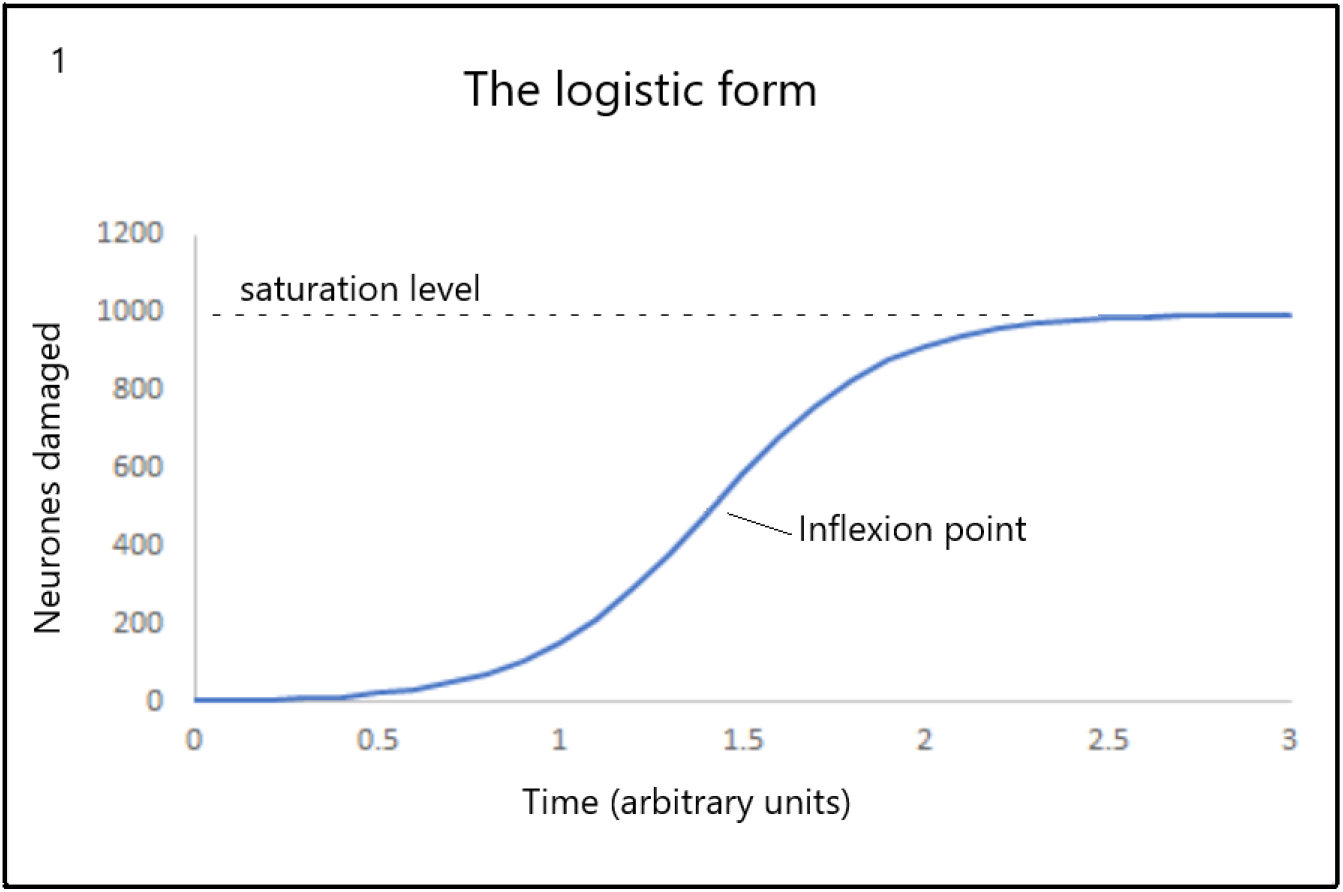
The classic logistic curve. The schematics used in this paper are copied from an accompanying Excel spreadsheet. Users may wish to explore the effects of changing the model variables. Model variables entered at “Main analysis” C11 to C14 in the accompanying Excel tool.

On the other hand, symptoms of NDD are often detectable before any candidate proteins are detectable, such proteins are not specific to any particular kind of NDD and thus far no medical interventions have had any detected^7^ effect on neurodegeneration despite having clear effects on target proteins.

The curve in Fig. 1 is explained as follows. Each infection-like event leads to the generation of further transmissive infective agents, leading to further infection-like events. At first an exponential increase in total infection sites with time is expected. As the number of uninfected sites reduces, the opportunity for each transmissive agent to cause a new infection reduces and the rate of new infection duly decreases. In the last 15% the logistic again closely resembles an exponential. In other words, in the last 15% the rate of loss of neurones decreases, with an exponential form, as more neurones are lost. At no other point is there a single exponential decay in the rate of neurone loss. It should be noted that if the logistic model is correct, the rate of neurodegeneration must increase as time passes provided less than half the neurones have been lost.

Eventually all sites (i.e., 1000 in Fig.1) have been infected.

Clinically, the frequency, duration and magnitude of symptom events increases until symptoms are more common than remission, eventually, becoming permanent. Just as an infection-like model would predict. It is common for logistic curves to be fitted to such clinical data^8^ (see Fig 1c Kang et al.).

On the other hand, the detailed study of neurone death rates shows only a reduction regardless of clinical status^9^. It is also noted that subjects who have been or soon will be diagnosed with one or more NDD have faster rates of neurone loss than those who remain undiagnosed^10^. But for a logistic, the rate must increase at first and only start to decrease when half the neurones have been lost. Instead, with or without NDD and with advanced NDD the same shape of steadily decreasing neurone death rate trajectory is obtained, but with different slopes. The shape of this trajectory is mathematically a single exponential decay. The rate of loss decreases as time progresses. It does not at any stage follow a logistic form. A single exponential decay should only be observed when more than 85% of neurones have been lost.

The conventional interpretation of clinical observation and actual neurone death rates are therefore seemingly incompatible.

The aim here is to show by means of schematics that the clinical logistic and the carefully observed single exponential are one and the same phenomenon.

An accompanying Excel document includes all of the results illustrated here and allows the user to experiment for themselves, within limits.

## 2. Axiomatic

Brain function is dependent on neurones and their connections. Loss of neurones threatens brain function. Neurones are insufficiently replaced in adulthood.

Therefore, if brain function was directly linked with neurone number a steady lifelong decrease of function would be predicted.

## 3. Threshold effects

In contrast with expectation, there are threshold effects^11,12,13^ in brain function.

The brain has the capacity to adapt to functional change associated with loss of neurones. Loss of neurones is compensated for by redundancy and degeneracy. With time, and sometimes very quickly, recruited neurones and alternative pathways become fully integrated and the affected brain function can be fully restored. The capacity to adapt is often described as reserve^14^ and is influenced by many factors including genetics, life experiences and education. So, while there is a clear general correlation of cognitive, memory and perceptual speed with neurone number the spread is very large (see Ref_11_). For a given cognitive score the neurone numbers can vary by multiples of two or four. The conclusion is that brain function and neurone number are related, but at least some of the time, they are not proportionate.

This adaptive capacity continues to cope very well, but not perfectly, with neurone loss until the effective supply of neurones available for recruitment/reorganisation is exhausted.

In other words, in its early stages, neurone loss may have a temporary effect on brain function, but later, as the adaptive capacity is exhausted, function loss becomes permanent. The situation is clearly captured in the following statement^15^ by S Przedborski and colleagues (with permission):

> *“Because, almost invariably, there is significant cellular redundancy in neuronal pathways, the onset of symptoms does not equate with the onset of the disease. Instead, the beginning of symptoms corresponds to a neurodegenerative stage at which the number of residual neurons in a given pathway falls below the number required to maintain normal functioning of the affected pathway*.*”*

Studies show that the proportion of neurone loss before the effect of functional loss is measurable and ranges from 10% in specialised olfactory functions to 80% in the substantia nigra. More typically, lasting and important functional loss occurs at between 30% and 50% of neurones lost^16^. The effect is described as a threshold (Fig. 2 red line).

**Fig. 2.**
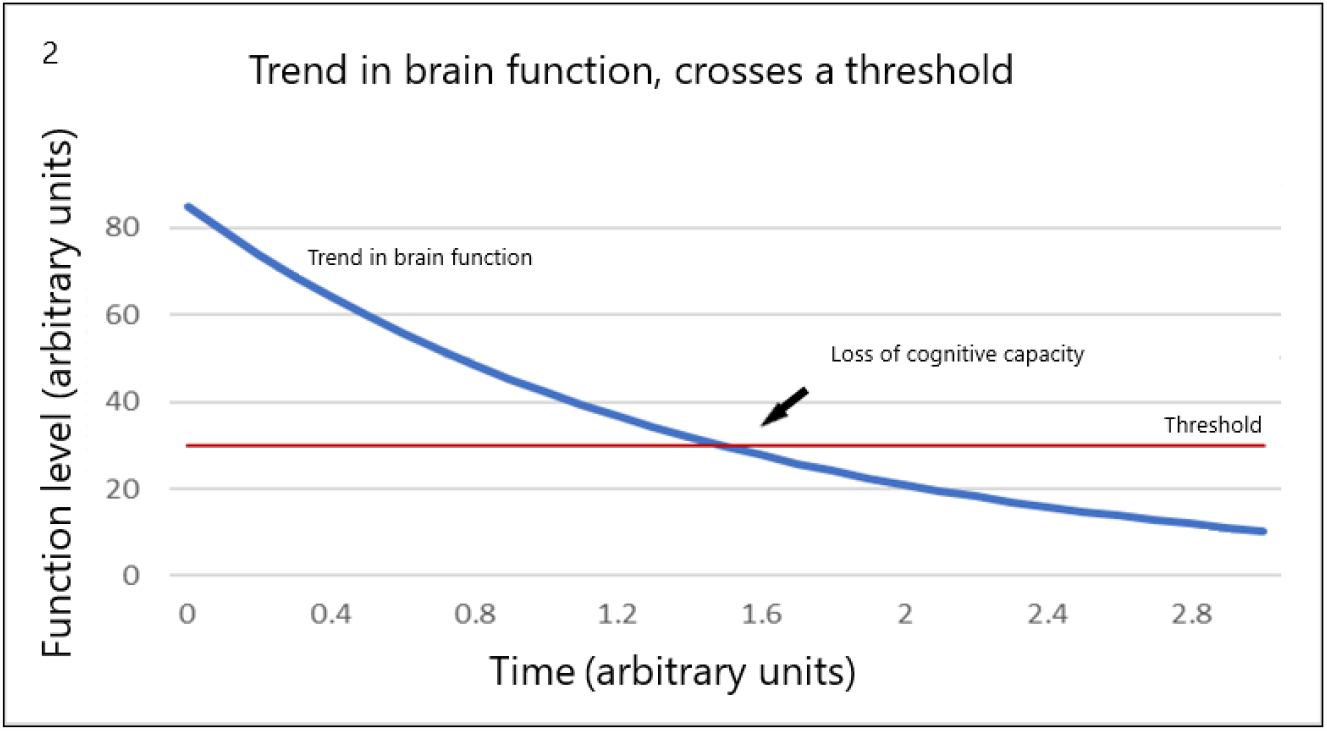
The blue curve represents the brain function effect of the advancing loss of neurones. The red line represents an effect threshold. Above threshold, the associated cognitive performance can be restored or partly restored. As the function level approaches and then falls below the threshold, the cognitive performance is more permanently impaired and then lost. Model variables entered at “Main analysis” E60 to E64 in the Excel tool.

Figure 2. represents the average effect of the observed exponential neurone loss curve (blue) and suggests that neurodegeneration only has lasting effects on cognitive performance once the threshold has been passed (red).

There are two direct results of the threshold-crossing model:

1. It is not that neurodegeneration migrates from one brain part to another it is just that the date of threshold-crossing varies from one part or set of parts to another. Time variation being explained by thresholds and rates of loss of neurones being different from one part to another. The associated effects e.g., short term memory and speech impairment, seem to ‘move around the brain’ but only because there is a sequence to threshold exceedance. Different brain functions effectively age at different rates^17^ and have different thresholds.
2. The repeated processes of loss and adaptation imply that the real effect of neurone loss on brain function and associated cognition is not a smooth process, but must fluctuate. The nature and timescales of the fluctuations must depend on the relationship between neurone number and function which could be localised or, more integrative. The mathematical form of these fluctuations is not known^18^, but once understood may be clinically informative. Cognitive fluctuations differ by type of dementia^19^. Fluctuations in functional network connectivity have been observed^20^ in people with and without dementia. The larger fluctuations being associated with the disease state. While each of these observations could indicate very different mechanisms it is proposed that each is related to changes in the level of function. If so, the point is that a static functional performance is not the norm.

## 4. Modelling the effect of brain function fluctuation

The simplest way to explore the effect of combing a trend in function level with function level fluctuations and a threshold is illustrated in Fig. 3 (below). Here the solid blue line from Fig. 2 has been replaced with a normally distributed fluctuation effect. A normal distribution would be more likely when there is a single localised mechanism of effect.

**Fig. 3.**
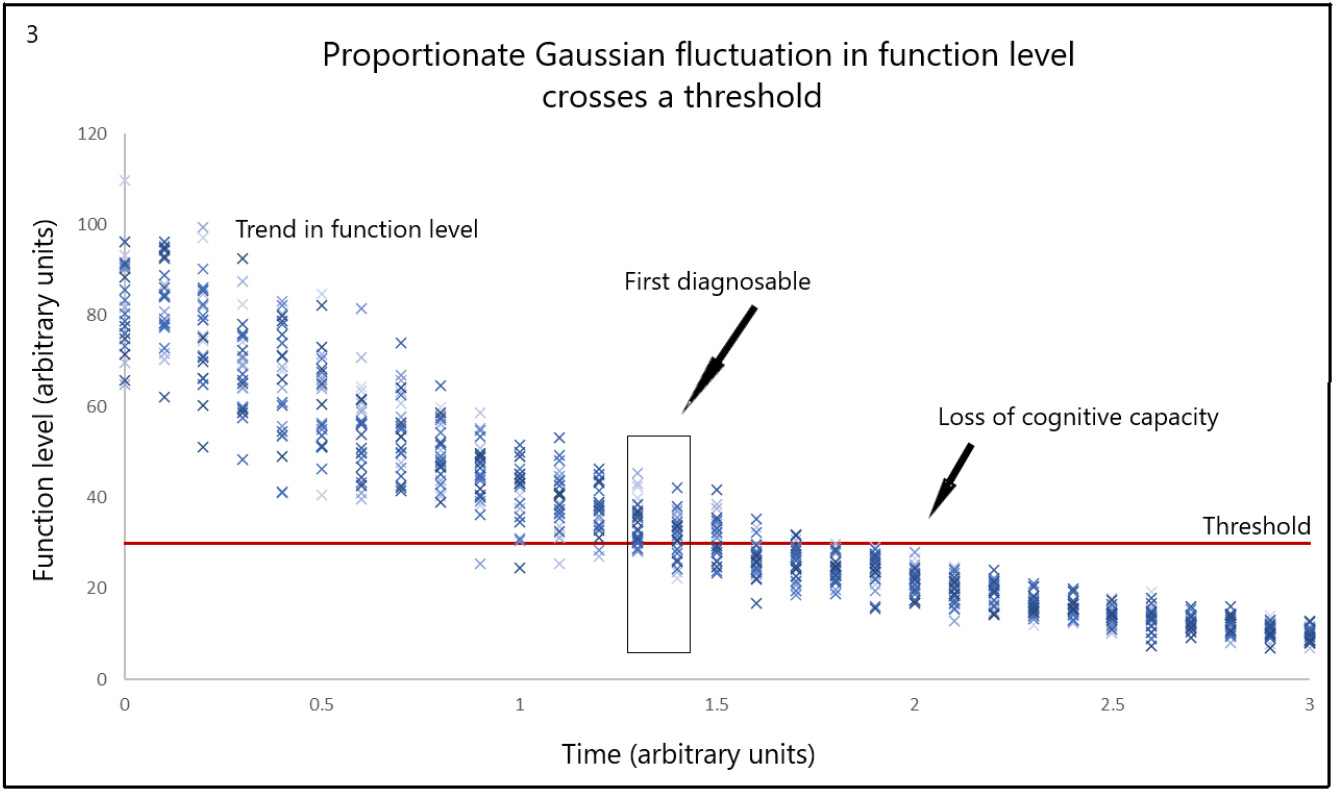
Functional performance fluctuates by a fixed proportion around a general trend driven by the gradual loss of neurones. The standard deviation was calculated as 21% of the function level. Trend variables are entered at “Main analysis” E60 to E64 and the % SDEV is entered at “Main analysis” R58 in the Excel tool.

From left to right:

1. Early in the function level trend there are very few occasions when fluctuations result in crossing the threshold.
2. The probability of threshold exceedance increases as time passes and further neurones are lost.
3. The case becomes diagnosable around the time range as indicated. This is calculated using the data plotted in Fig. 14.
4. Despite fluctuations, there is a point where the threshold is effectively out of reach and the use of that function is permanently lost.

The model accords with the case history of neurodegenerative disease.

The probability of threshold exceedance in this model can be calculated and plotted against time (Fig. 4). This is practically demonstrated in the accompanying Excel document at “Main analysis” figure at cell AG60.

**Fig. 4.**
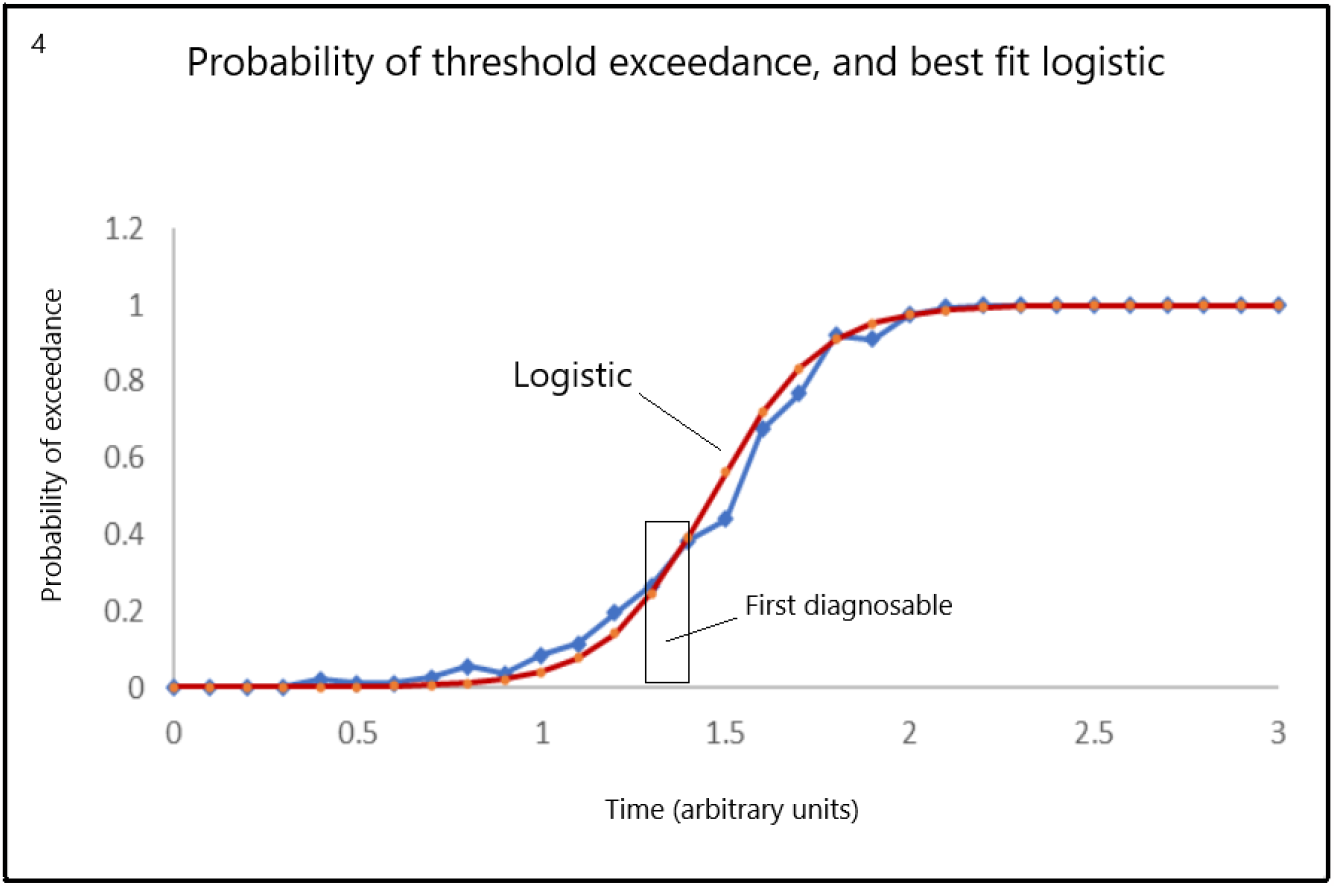
Using the model depicted in Fig 3. The probability that the function threshold is exceeded is calculated (blue points). The probability of exceedance is reasonably well described by a fited logistic (red curve). The region where a person would be first diagnosable is indicated and derived from the data plotted in Fig 14. The above figure is located at “Main analysis” at cell AG60 in the accompanying Excel tool.

Given that the relationship between effective function level and fluctuation is not known it is helpful to test a wider range of %SDEV values. Fig. 5 shows the effect of an order of magnitude change in %SDEV values on the generation of a logistic. In each case, a logistic is generated.

**Fig. 5.**
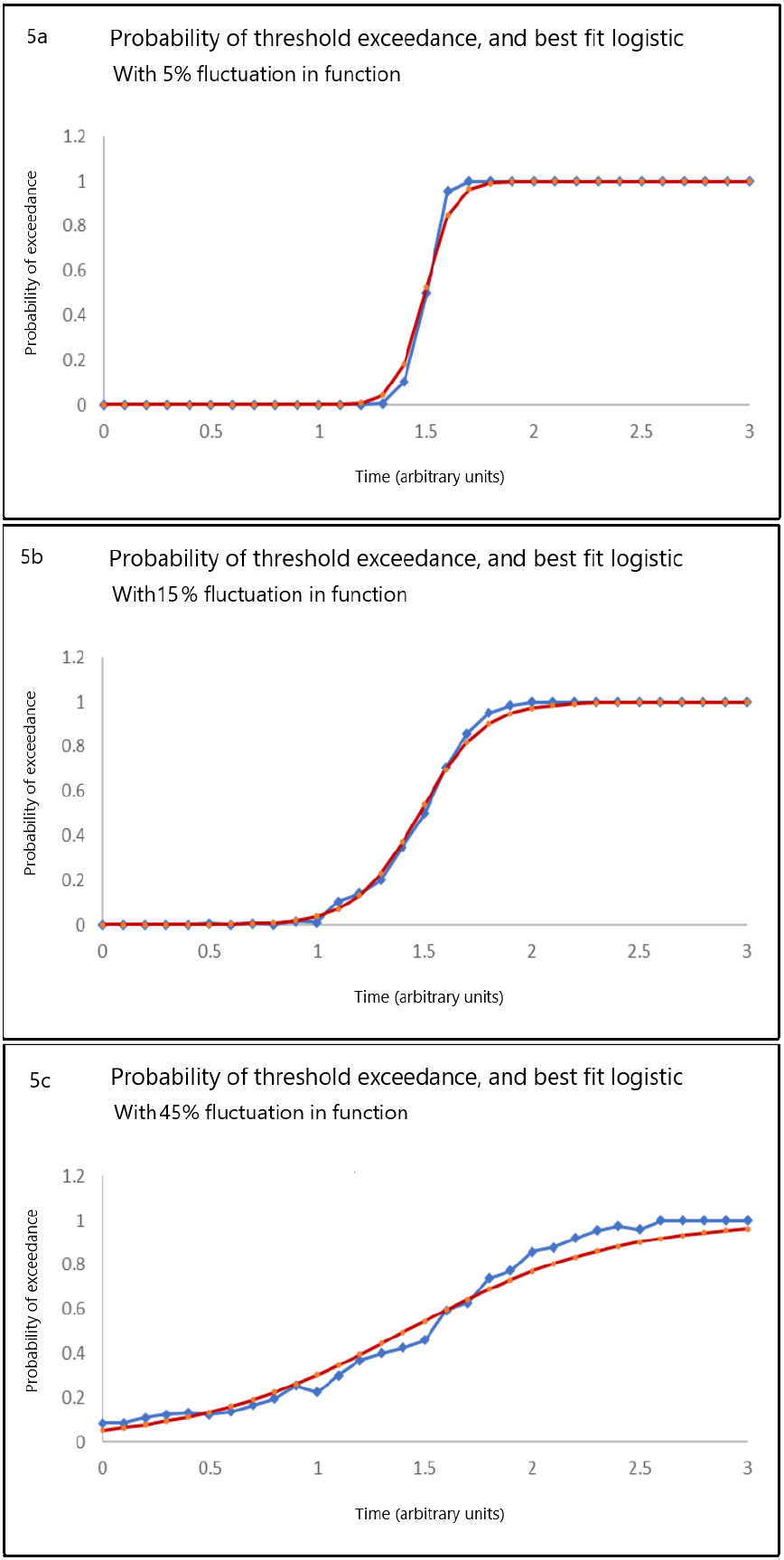
The probability of threshold exceedance (blue points) using 5%, 15% and 50% of function level as the SDEV value for normally distributed fluctuations in function level, a, b and c respectively. In each case the same function level trend and threshold are adopted. In each case the resulting probability of exceedance curve is reasonably well described by a fitted logistic (red curve). The % SDEV value is entered at “Main analysis” R58 in the Excel tool.

The time scale on the x axis is arbitrary but clinical experience suggests it corresponds to between 5 and 15 years. The probability of exceedance curve follows the observed trend in frequency of cognitive decrements.

A simple exponential decay in neurone number, which leads to an advancing but fluctuating loss of functional performance, combined with a functional threshold generates the clinically observed logistic or sigmoid form with no need at all for an infection-like mechanism.

## 5. Is the generation of a sigmoid function a generalisable result?

Generalisability is addressed in a series of questions:

Q1. Would the result in Fig. 4. occur if the amplitude of fluctuations was constant?

Q2. What if the threshold also has fluctuations?

Q3. What if the relationship between neurone loss and functional performance resulted in a linear instead of an exponential decline?

Q4. Would a lognormal distribution of fluctuations lead to a logistic trend?

Q5. Would random fluctuations in functional performance lead to a logistic trend?

Q6. Does it matter from which direction the threshold is crossed?

The accompanying Excel spreadsheet works through these questions and in each case, an approximately logistic form is generated. While this is not a mathematical proof of generalisability this is clearly credible and is easy to grasp by the non-mathematician.

The following Figures are copied from the spreadsheet.

Q1: Constant amplitude fluctuation. This might occur when the cause of loss of function is remote from the function centre.

**Fig. 6.**
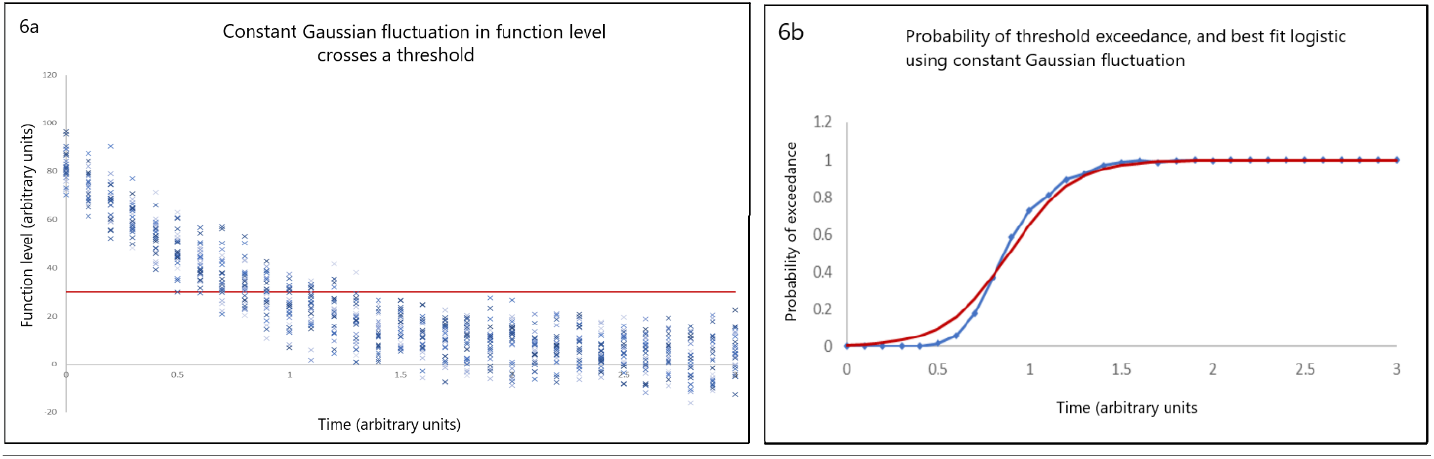
On the left (6a) functional level fluctuates at a constant amplitude. On the right, (6b) the probability of threshold exceedance (blue dots) is reasonably represented by a fitted logistic (red curve). Trend variables are entered at “Main analysis” E107 to E111 and the % SDEV is entered at “Main analysis” R105 in the Excel tool.

Q2: Fluctuation in both the functional performance and in the threshold.

**Fig. 7.**
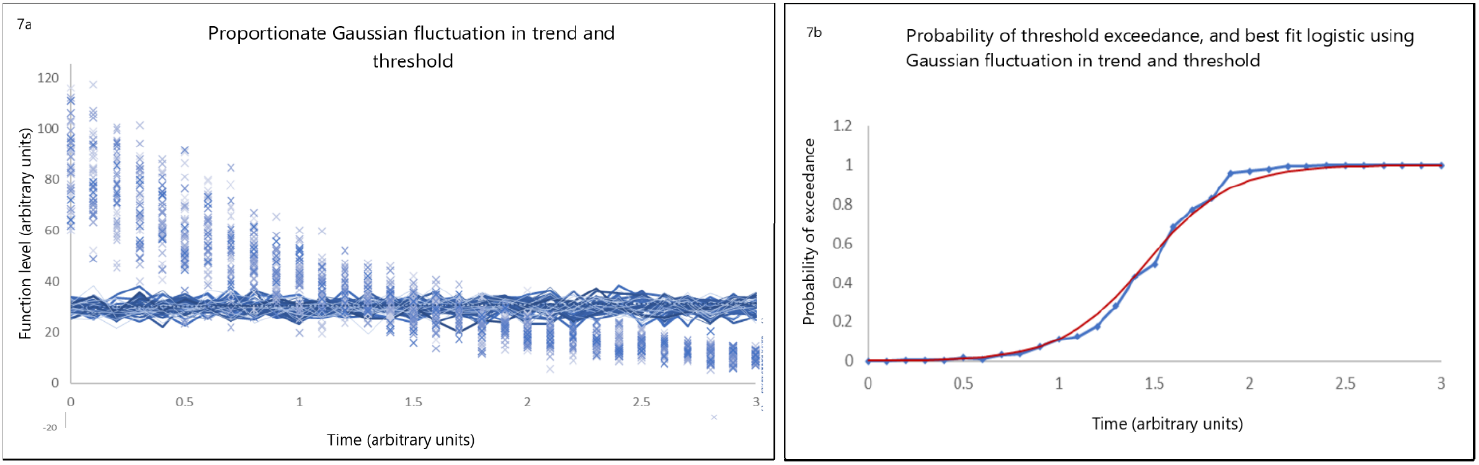
On the left (7a) functional level and threshold both fluctuate with a proportionate amplitude. On the right, (7b) the probability of threshold exceedance (blue dots) is reasonably represented by a logistic (red line). Trend variables are entered at “Main analysis” E148 to E152 and the % SDEV for trend and threshold are entered at “Main analysis” R150 and R151 respectively, in the Excel tool.

Q3: Linear decay, fluctuations in both.

**Fig. 8.**
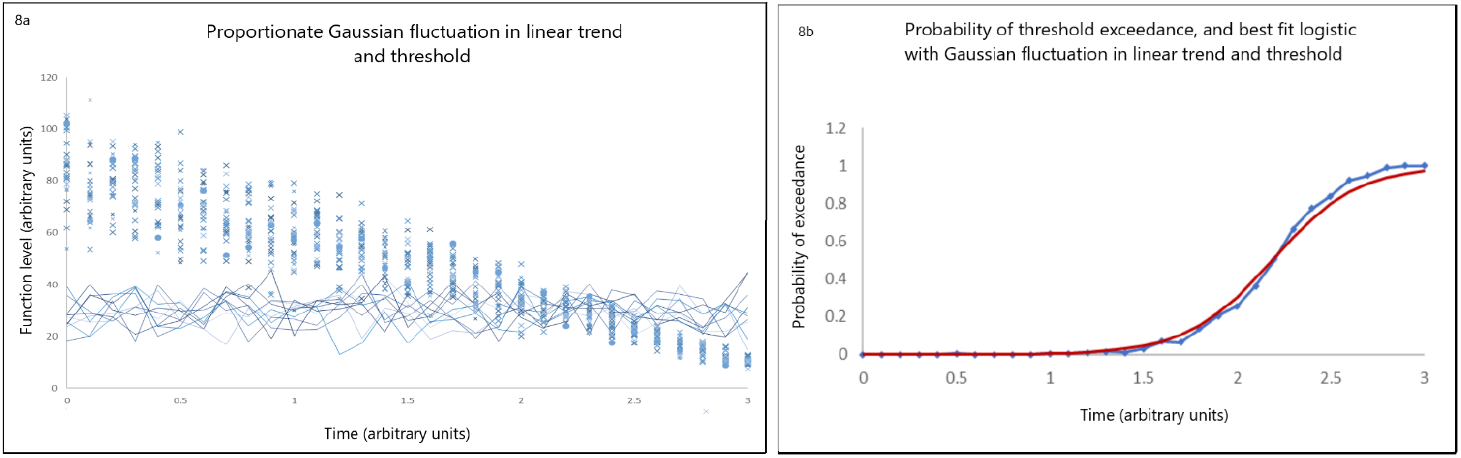
On the left (8a) the functional level follows a linear trend with proportionate fluctuation. On the right (8b), the probability of threshold exceedance (blue dots) is reasonably represented by a logistic (red line). Trend variables are entered at “Main analysis” E189 to E193 and the % SDEV for trend and threshold are entered at “Main analysis” N186 and N187 respectively, in the Excel tool.

Q4: Lognormal fluctuations in functional performance and in threshold. As might be expected when several independent mechanisms are acting.

**Fig. 9.**
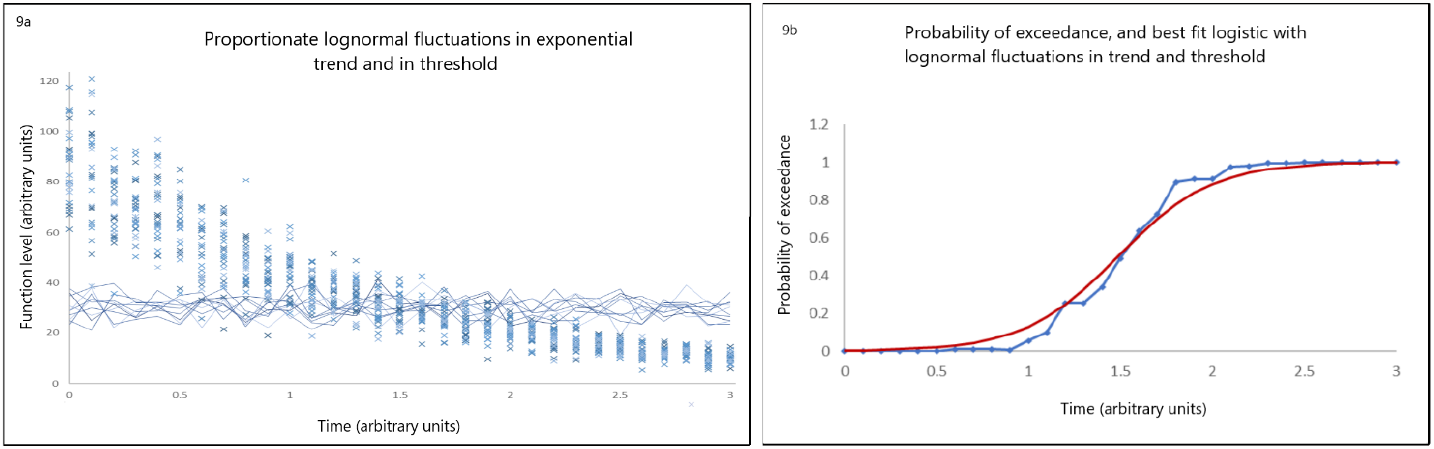
On the left (9a), the functional level follows the exponential trend with fluctuations distributed lognormally. On the right (9b), the probability of threshold exceedance (blue dots) is reasonably represented by a logistic (red line). Trend variables are entered at “Main analysis” E226 to E230 and the % SDEV for trend and threshold are entered at “Main analysis” E232 and E234 respectively, in the Excel tool.

Q5: Random fluctuations in functional performance and threshold. That is, all fluctuations equally likely; a very unlikely scenario in biology.

**Fig. 10.**
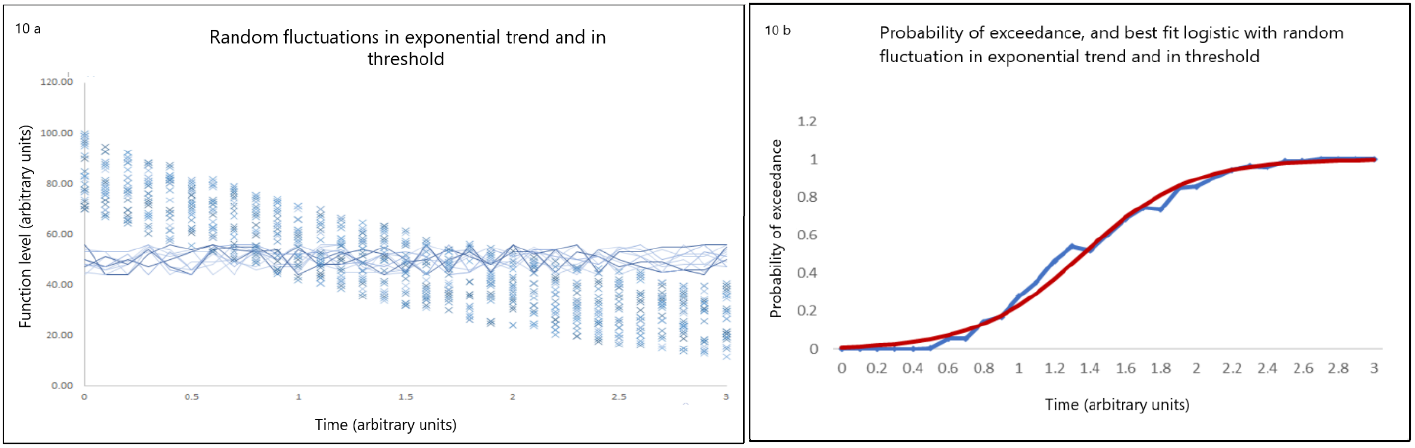
On the left (10a), functional level and threshold both experience random fluctuations. On the right (10b), the probability of threshold exceedance (blue line) is reasonably represented by a logistic (red line). Trend variables are entered at “Main analysis” E305 to E309 and the % SDEV for trend and threshold are entered at “Main analysis” E313 and E315 respectively, in the Excel tool.

While this is not an exhaustive analysis the results in Fig. 4. to Fig. 10. suggest confidence that when brain functions approach, cross and then exceed a threshold, and do so with fluctuations, the probability of exceedance follows a logistical or sigmoid form.

Q6: Crossing a threshold from below. It makes no difference if the threshold is approached from above, or from below:

**Fig. 11.**
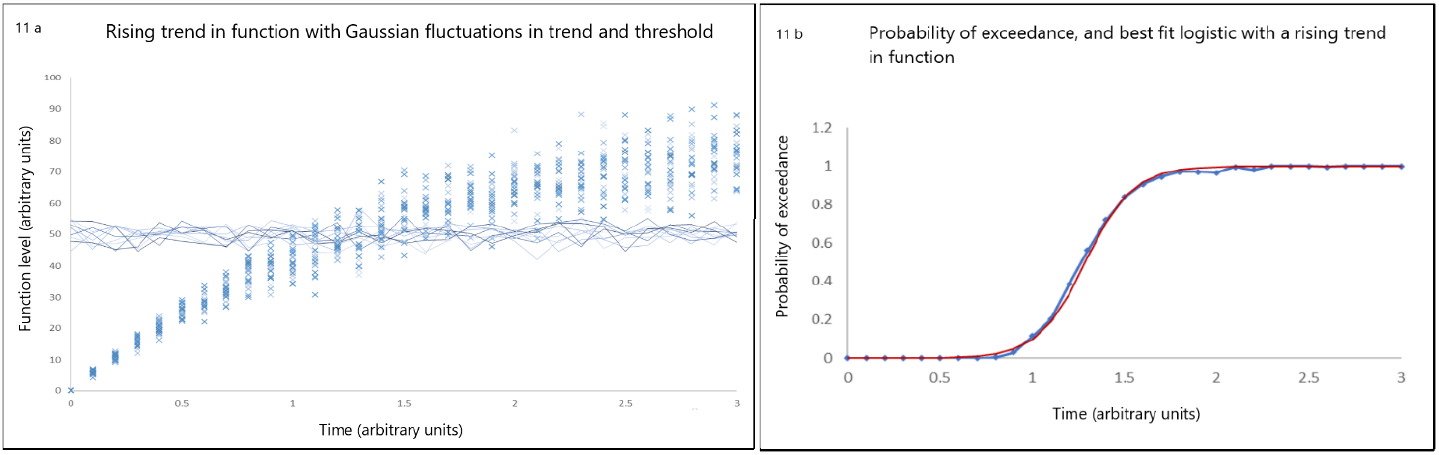
On the left (11a), functional level increases, and there is proportionate fluctuation in level and threshold. On the right (11b), the probability of exceedance (blue dots) is reasonably represented by a logistic (red line). Trend variables are entered at “Main analysis” E265 to E269 and the % SDEV for trend and threshold are entered at “Main analysis” E273 and E275 respectively, in the Excel tool.

## 6. Observable cognitive capacity

Having argued that the sigmoid form in frequency of cognitive deficits can be explained by a threshold-crossing model the question is whether the same model can explain trends in measured cognitive performance.

Given the adaptive capability of brain functions and provided the functional performance is well above threshold then the measured level of cognitive capacity will on average be steady or decline slightly if adaptation is not perfect. At the other extreme, once permanently below threshold the cognitive capacity is lost completely.

A model is needed to link the two extremes. While it is clear that a cognitive capability does not go instantly from normal to completely lost, the transition between normal and lost performances may be related to the probability of exceedance and to the height of the threshold relative to the present function level. In some way, these parameters may measure the strain under which the adaptive process is placed. It is proposed here that on close approach to threshold, adaptation is increasingly under strain and the completeness of adaptation is reduced. Incomplete adaptation leads to reduced cognitive capability.

Normalised performance may be modelled by:

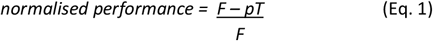

Where *F* is the degree of function, *p* is the probability of exceeding the threshold and *T* is the threshold. Using the example of 20% proportionate Gaussian fluctuations in *F* and 10% proportionate Gaussian fluctuations in *T* and where *F* approaches *T* as a single exponential we get, for an individual:

Experimental data for groups of people will give a spread of starting points on the y axis unless the data is first normalised. Each person would also have a different curvature and each a different date of diagnosis or some other reference date. Despite this, there may well be groups of people with similar curvature and this will be apparent if the time origin is chosen to superimpose dates of diagnosis.

If an additional strain factor *K* is added, perhaps representing some metabolic strain, normalised performance could be represented as:

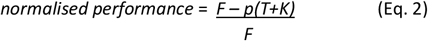

With a positive *K* the slope of the decline in performance is faster as represented in Fig. 13.

**Fig. 14.**
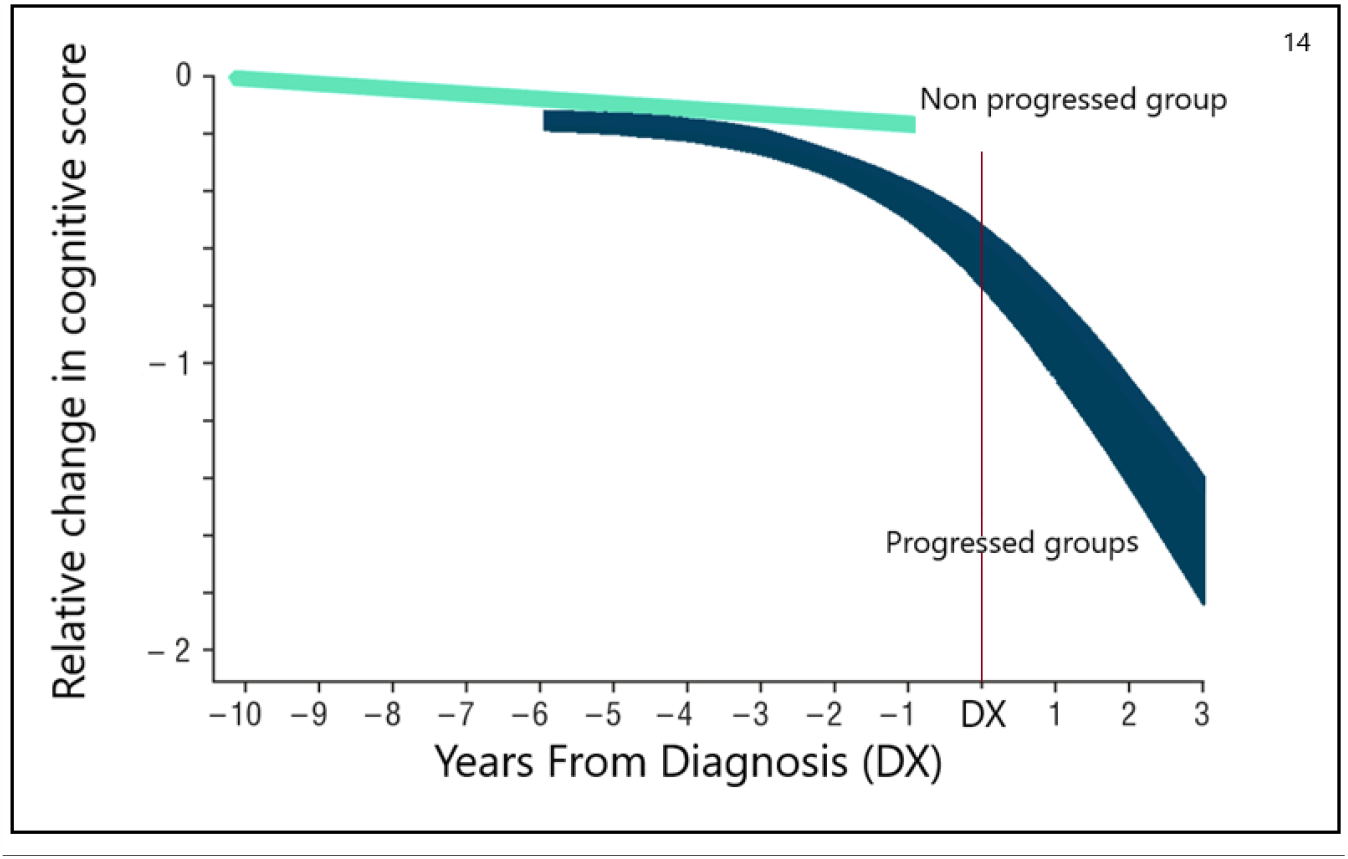
Time evolution of cognitive test scores as found in well-conducted research. The reference date is the date of diagnosis (DX). The distinction between progressed and non-progressed groups is in how close to threshold they are.

In Figures 12 and 13 there is no obvious point of inflexion at the half height of the curves. Given the probabilistic nature there could sometimes appear to be a slightly extended tail.

**Fig. 12.**
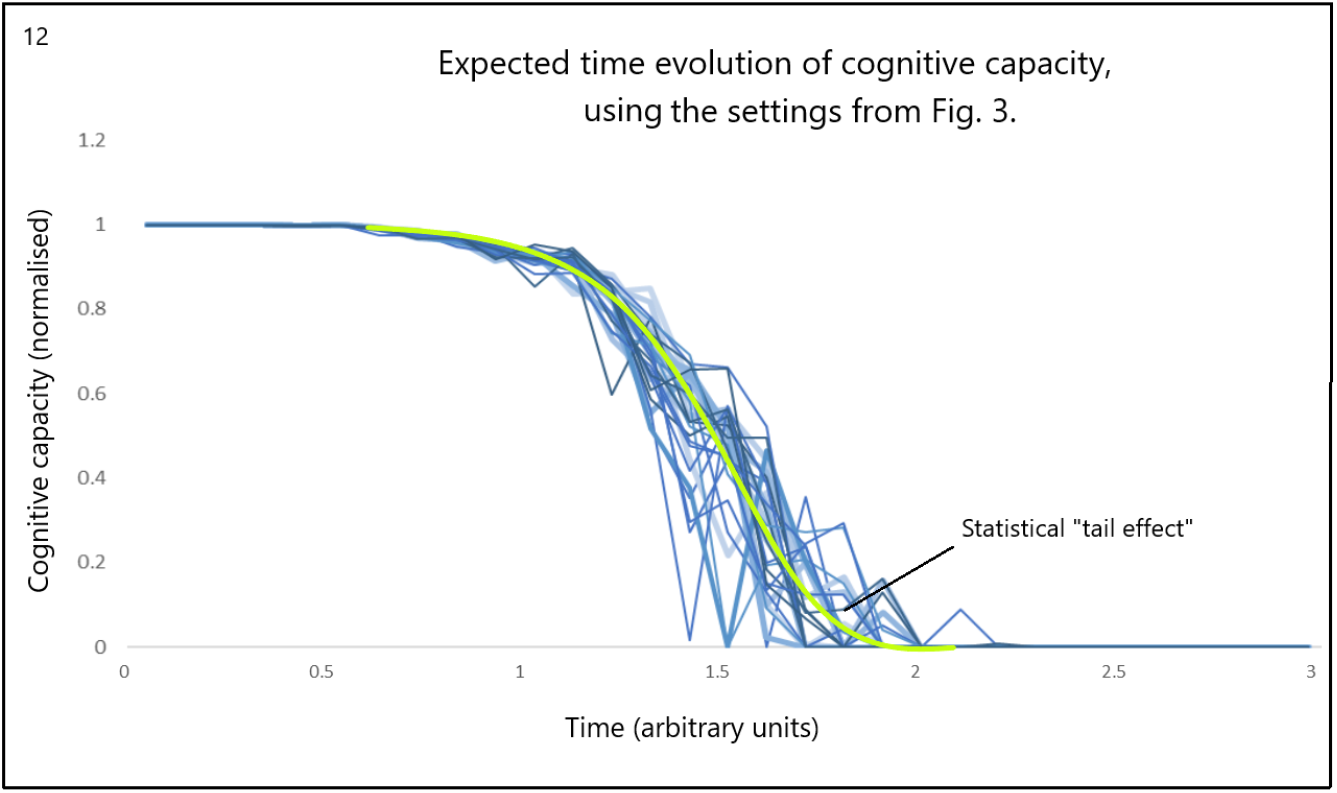
Time evolution of cognitive capacity as predicted by a simple threshold-crossing model as expressed in equation 1. There is no inflexion at half height but a tail effect may be observed. Trend variables are entered at “Main analysis” E148 to E152 and the % SDEV for trend and threshold are entered at “Main analysis” R150 and R151 respectively, in the Excel tool. The figure is at “Main analysis” V351.

**Fig. 13.**
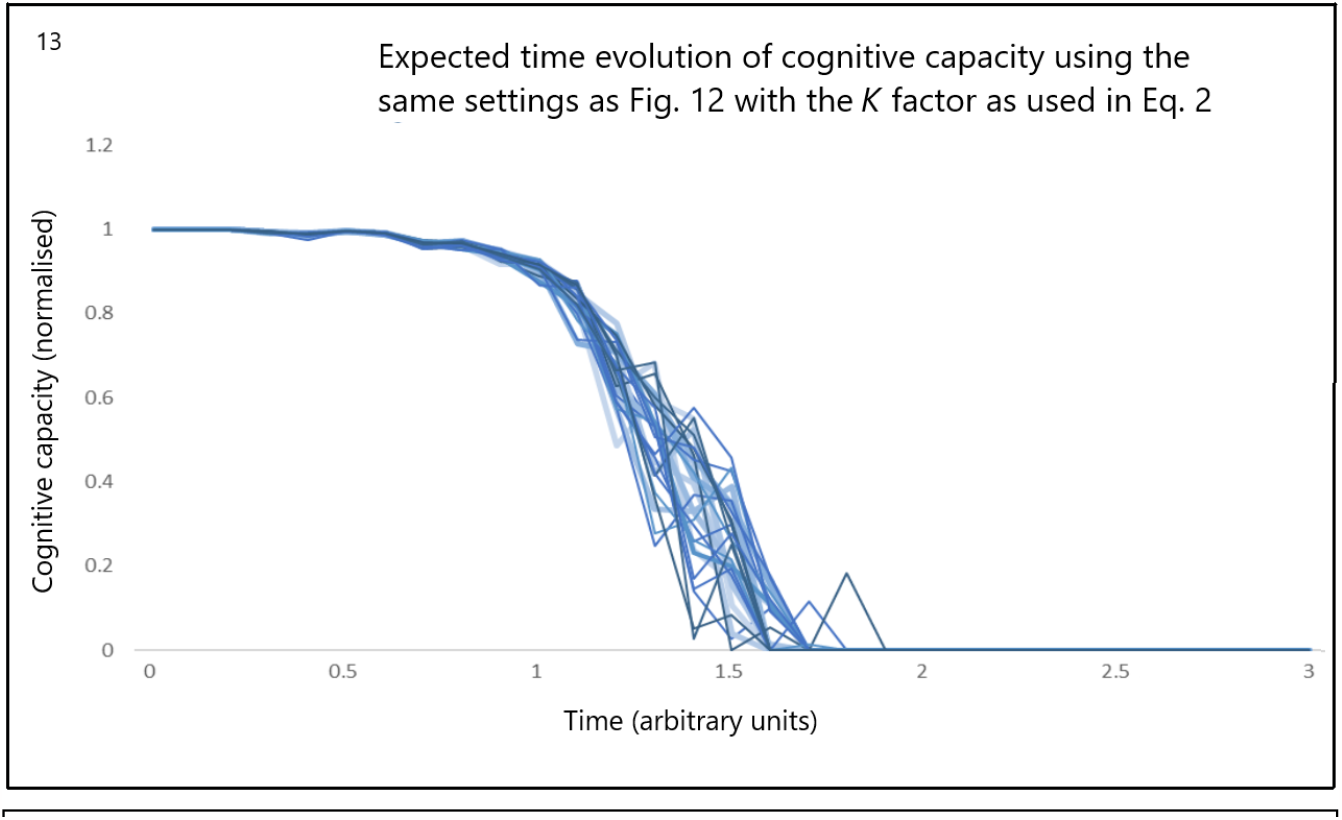
Time evolution of cognitive capacity as predicted by a simple threshold model with an additional strain factor as expressed in equation 2. With a positive *K*, the decline in cognitive capacity is faster. Variables are entered as in Fig. 12 but the *K* factor is entered at “Main analysis” X350.

Could these predicted effects be found in clinical data? In designing longitudinal studies of cognitive performance care must be taken not to combine data from groups of people with different outcome status. There is no reason to suppose that the rate of loss in normal controls is now or ever will be the same as in mild cognitive impairment or diagnosed disease. However, this homogeneity assumption is the norm^21,22^. What is known is that the rate of loss cognitive performance is slowest in normal controls, and fastest in those diagnosed^23^. Likewise calendar age is not a useful x axis, but years before and after diagnosis, or reference date, would be better. Some studies combine the homogeneity assumption and calendar age assumption simultaneously^24^, making them very hard to interpret as a result.

As a general observation, the imprecision evident in the longitudinal measurement of cognitive performance suggests that studies spanning less than 5 years are unlikely to yield meaningful longitudinal trajectories. Such studies would not be useful in assessing the predictions made using equations 1 and 2.

The trend identified in Fig 12 is an excellent match with data from well-conducted longitudinal studies^22,25,26,^. Figure 14. represents the form of data found in well-conducted studies:

Acceleration of cognitive decline with level of that decline is well-established^27^. However, not all studies^28^ find exactly these trends.

## 7. Implications

If the observed logistic was indeed a result of a transmissive replicatable agent then its identification could lead to a medical intervention, e.g., a replication inhibitor.

However, neurodegeneration is resolutely of a single exponential decay form, long before the last 15% of neurones are all that remain, and there are thresholds above which functional restoration is possible. When combined, these would likely result in a logistic or sigmoid form of temporal clinical development.

A single exponential trend means there is no concerted^29^ effect in neurodegeneration, just independent neurone death events. An infection-like model could not explain this result.

The threshold-crossing model hypothesises that those people who develop clinical NDD do so simply because their brain ages fast enough to generate symptoms before death. This is either because their neurodegeneration just happens to be faster or the thresholds are higher, or both.

The threshold-crossing model also explains why NDD seems to move around the brain. Short-term memory and reasoning for example, have different thresholds and ageing rates. Rather than movement of the site of neurone loss leading to specific function loss the clinical signs indicate the order in which the threshold is crossed. There is no reason to assume that in the absence of some disease agent all brain functions age at the same rate. In short, there is no need to invoke a transmissive infection-like agent.

In addition to searching for a transmissive replicative neurodegenerative agent the focus of research could usefully shift to understanding why some people’s brains age more quickly than others, why some functions age more quickly and what determines the function threshold. Interventions in any of these could spare many people from NDD. Perhaps the removal of amyloid beta plaques doesn’t affect neurodegeneration but simply allows neurones to be more metabolically active and thereby be more functionally resilient.

Why is it that childhood adversity increases the risk of detection of NDD? Why does intellectual stimulation reduce the risk? A strong possibility is that the answer rests in the threshold-crossing model rather than in some, as yet, elusive transmissive replicative neurogenerative agent present in people with NDD.

More generally, a sigmoid response function can be generated by many different mathematical forms, including the logistic. Godeau *et al*. ^30^ identify twelve mathematical formulae that generate a sigmoid curve. Each is based on a different biological relationship between the variables on the x and y axes (respectively the explanatory and explained variables). What these relationships have in common are lower and upper asymptotes, a continuous positive slope from lower to upper asymptote, a point of inflexion, usually around half-way between the asymptotes, and a degree of rotational symmetry about the inflexion point. Though not precise, the sigmoid curves generated by the probability of threshold exceedance are better thought of as cumulative Gaussians. The question then is whether biological data is precise enough to be certain of distinguishing between the different mathematical forms and therefore the underlying biological relationships? In my view, the answer is, not yet.

The threshold-crossing model of loss of function is not new, but to the best of my knowledge there has been no literature showing that this model leads to the observed sigmoid NDD clinical trajectory in frequency or to the observed trend in cognitive performance.

## 8. Conclusion

The threshold-crossing model provides a consistent explanation of the phenomenology of neurodegenerative disease. The infection-like model does not.

## 9. Acknowledgements

My thanks to Professor P Langford and Dr C Serpell for their critical reading of this manuscript in draft form.

## Excel Tool

An Excel tool was used to test the threshold-crossing model. A copy is available from Dr Andrew Auty upon request.

The tool requires macros to be enabled and it may not work on older versions of MS Office. These problems are readily solved by university IT departments.

## References

1 McKendrick AG, Pai M . Xl V.—The Rate of Multiplication of Micro-organisms: A Mathematical Study. Proceedings of the Royal Society of Edinburgh. (1912) 31:649–653. doi:10.1017/S0370164600025426

2 Lewis PA, Spillane JE (Eds). Alzheimer’s Disease and Dementia. In, The Molecular and Clinical Pathology of Neurodegenerative Disease. Chapter 2 p 25-82. (2018) Academic Press. ISBN 978-0-12-811069-0.

3 Mudher A, Colin M, Dujardin S, Medina M et al. What is the evidence that tau pathology spreads through prion-like propagation? Acta Neuropathol Commun. 2017;5(1):99. doi: 10.1186/s40478-017-0488-7.

4 Stopschinski BE, Diamond MI. The prion model for progression and diversity of neurodegenerative diseases. Lancet Neurol. 2017;16(4):323–332. doi: 10.1016/S1474-4422(17)30037-6.

5 Wilson RS, Capuano AW, Bennet DA, Schneider JA, Boyle PA. Temporal course of neurodegenerative effects on cognition in old age. Neuropsychology. 2016;30(5):591–9. doi: 10.1037/neu0000282.

6 Braak H, Braak E. Neuropathological stageing of Alzheimer-related changes. Acta Neuropathol. 1991;82(4):239–59. doi: 10.1007/BF00308809.

7 https://www.alz.org/alzheimers-demena/treatments/aducanumab Accessed on 14th November 2023.

8 Kang K, Dumitrescu L, Mukherjee S, Lee ML et al. Synchronized sigmoidal mixed-effects model for dynamics of cognitive decline relative to onset of Alzheimer’s disease in aging adults in the Alzheimer’s Disease Neuroimaging Initiative (ADNI) study. Alzheimer’s and Dementia. (2022) 18 Supp11: e063496.

9 Gómez-Isla T, Hollister R, West H, Mui S, et al. Neuronal loss correlates with but exceeds neurofibrillary tangles in Alzheimer’s disease. Ann Neurol. 1997;41(1):17–24. doi: 10.1002/ana.410410106.

10 Sabuncu MR, Desikan RS, Sepulcre J, Yeo BT et al. Alzheimer’s Disease Neuroimaging Initiative. The dynamics of cortical and hippocampal atrophy in Alzheimer disease. Arch Neurol. 2011;68(8):1040–8. doi: 10.1001/archneurol.2011.167.

11 Welsh-Bohmer KA. Defining “prodromal” Alzheimer’s disease, frontotemporal dementia, and Lewy body dementia: are we there yet? Neuropsychol Rev. 2008;18(1):70–2. doi: 10.1007/s11065-008-9057-y.

12 Kelly SC, He B, Perez SE, Ginsberg SD et al. Locus coeruleus cellular and molecular pathology during the progression of Alzheimer’s disease. Acta Neuropathol Commun. 2017;5(1):8. doi: 10.1186/s40478-017-0411-2.

13 Brown TP, Rumsby PC, Capleton AC, Rushton L et al. Pesticides and Parkinson’s disease--is there a link? Environ Health Perspect. 2006;114(2):156–64. doi: 10.1289/ehp.8095.

14 Stern Y, Arenaza-Urquijo EM, Bartrés-Faz D, Belleville S et al. Whitepaper: Defining and investigating cognitive reserve, brain reserve, and brain maintenance. Alzheimer’s Dement. 2020;16(9):1305–1311.

15 Przedborski S, Vila M, Jackson-Lewis V. Series Introduction: Neurodegeneration: What is it and where are we? J Clin Invest. 2003;111(1): 3–10.

16 Riddle DR, (Ed). Design-Based Stereology in Brain Aging Research. In, Brain Aging: Models, Methods, and Mechanisms. Chapter 4. P. 64-88. Boca Raton (FL): CRC Press/Taylor & Francis; 2007.

17 Dehkordi SK, Walker J, Sah E, Bennet E, et al. Profiling senescent cells in human brains reveals neurons with CDKN2D/p19 and tau neuropathology. Nat Aging. 2021;1(12):1107–1116. doi: 10.1038/s43587-021-00142-3.

18 Papo D. Functional significance of complex fluctuations in brain activity: from resting state to cognitive neuroscience. Front Syst Neurosci. 2014;8:112. doi: 10.3389/fnsys.2014.00112.

19 Van Dyk K, Towns S, Tatarina O, Yeung P et al. Assessing Fluctuating Cognition in Dementia Diagnosis: Interrater Reliability of the Clinician Assessment of Fluctuation. Am J Alzheimers Dis Other Demen. 2016;31(2):137–43. doi: 10.1177/1533317515603359.

20 Moguilner S, García AM, Perl YS, Tagliazucchi E et al. Dynamic brain fluctuations outperform connectivity measures and mirror pathophysiological profiles across dementia subtypes: A multicenter study. Neuroimage. 2021;225:117522. doi: 10.1016/j.neuroimage.2020.117522.

21 Kang K, Dumitrescu L, Mukherjee S, Lee ML et al. Synchronized sigmoidal mixed-effects model for dynamics of cognitive decline relative to onset of Alzheimer’s disease in aging adults in the Alzheimer’s Disease Neuroimaging Initiative (ADNI) study. Alzheimer’s Dement. 2022;18 Supp11: e063496.

22 Zhuo J, Zhang Y, Liu Y, Liu B et al. Alzheimer’s Disease Neuroimaging Initiative. New Trajectory of Clinical and Biomarker Changes in Sporadic Alzheimer’s Disease. Cereb Cortex. 2021;31(7):3363–3373. doi: 10.1093/cercor/bhab017.

23 Wang HF, Shen XN, Li JQ, Suckling J et al. Alzheimer’s Disease Neuroimaging Initiative. Clinical and biomarker trajectories in sporadic Alzheimer’s disease: A longitudinal study. Alzheimer’s Dement (Amst). 2020;12(1):e12095. doi: 10.1002/dad2.12095. https://alz-journals.onlinelibrary.wiley.com/doi/epdf/10.1002/dad2.12095.

24 A Pai, S Sommer, LL Raket, L Sørensen et al. Do Dementia Biomarkers Have A Sigmoid Trajectory? Insights From Non-Linear Mixed Effects Modeling. Alzheimer’s Dement. 2016;12(7):356. doi.org/10.1016/j.jalz.2016.06.659

25 Johnson DK, Storandt M, Morris JC, Galvin JE. Longitudinal study of the transition from healthy aging to Alzheimer disease. Arch Neurol. 2009;66(10):1254–9. doi: 10.1001/archneurol.2009.158. https://jamanetwork.com/journals/jamaneurology/article-abstract/798199

26 Kiddle SJ, Parodi A, Johnston C, Wallace C et al. Heterogeneity of cognitive decline in dementia: a failed attempt to take into account variable time-zero severity. bioRχiv. 2017: doi.org/10.1101/060830 .

27 Teri L, McCurry SM, Edland SD, Kukull WA et al. Cognitive decline in Alzheimer’s disease: a longitudinal investigation of risk factors for accelerated decline. J Gerontol A Biol Sci Med Sci. 1995;50A(1):M49–55. doi: 10.1093/gerona/50a.1.m49. https://academic.oup.com/biomedgerontology/article/50A/1/M49/616783

28 Boyle PA, Yang J, Yu L, Leurgans SE et al. Varied effects of age-related neuropathologies on the trajectory of late life cognitive decline. Brain. 2017;140(3):804–812. doi: 10.1093/brain/aww341.https://academic.oup.com/brain/article/140/3/804/2898323?login=false

29 https://mathworld.wolfram.com/ExponenalDecay.html Accessed on 15th Sept 2023.

30 Godeau U, Bouget C, Piffady J, Pozzi T et al. Lack of definition of mathematical terms in ecology: The case of the sigmoid class of functions in macro-ecology. Ecol Evol. 2020;10(24):14209–14220. doi: 10.1002/ece3.7016.

